# Genome-Wide Association Study of Ionomic Traits on Diverse Soybean Populations from Germplasm Collections

**DOI:** 10.1101/079731

**Authors:** Greg Ziegler, Randall Nelson, Stephanie Granada, Hari B. Krishnan, Jason D. Gillman, Ivan Baxter

## Abstract

The elemental content of a soybean seed is a determined by both genetic and environmental factors and is an important component of its nutritional value. The elemental content is chemically stable, making the samples stored in germplasm repositories an intriguing source of experimental material. To test the efficacy of using samples from germplasm banks for gene discovery, we analyzed the elemental profile of seeds from 1653 lines in the USDA Soybean Germplasm Collection. We observed large differences in the elemental profiles based on where the lines were grown, which lead us to break up the genetic analysis into multiple small experiments. Despite these challenges, we were able to identify candidate SNPs controlling elemental accumulation as well as lines with extreme elemental accumulation phenotypes. Our results suggest that elemental analysis of germplasm samples can identify SNPs in linkage disequilibrium to genes, which can be leveraged to assist in crop improvement efforts.

## Introduction

One of the biggest challenges facing agricultural research today is finding ways to improve crop yield and nutrition while farming in increasingly erratic climates and on more marginal lands. Throughout modern agriculture, crops have been bred for maximal yield under optimal environmental conditions. Farming marginal soils with insufficient fertilization or irrigation leads to dramatic decreases in crop yield. In addition, plants grown on marginal soils may exhibit a reduced nutritional profile, which is an important consideration for staple crops. To properly address these issues, we need to develop a more complete understanding of the genetic mechanisms underlying a plant’s response to various environmental stresses (Baxter and Dilkes 2012).

An important aspect underlying a plant's response to environmental stresses is its ability to regulate mineral nutrients. Apart from carbon and oxygen, a plant relies entirely on the bioavailable nutrients in the soil in which it is growing for survival. Soil nutrient bioavailability can vary drastically, not just as a result of soil composition, but also as a side effect of drought and flood conditions, changes in soil pH, and changes in the soil microbiome (FAO 1996). Understanding the uptake, regulation, transport, and storage of mineral nutrients under a variety of environmental conditions is essential to deciphering the complex relationship between a plant and its environment.

Single-seed ionomic profiles have proven both highly heritable and susceptible to environmental perturbations in maize (Baxter *et al.* 2014). This makes the study of the seed ionome a powerful tool for matching a plant’s genetic characteristics with its response to environmental perturbations. Both environmental and genetic properties can effect multiple elements in combination, resulting in genetic loci that might control different elements in different environments (Baxter 2015; Asaro *et al.* 2016). Additionally, once collected, apart from the possibility of external contamination, the elemental content of a seed sample is fixed. Tissue for ionomic analysis doesn't need to be specially stored or quickly analyzed after collection. Conveniently, this allows for the ionomic analysis of excess tissue collected for other purposes, without the necessity of a separate field experiment. Here we demonstrate the utility of leveraging existing germplasm by performing a genome-wide association study on ionomic traits in seed tissue measured from diverse soybean lines selected from the USDA Soybean Germplasm Collection.

## Results

### Experimental Design

The mission of the USDA-ARS National Plant Germplasm System (NPGS) is “to acquire, evaluate, preserve and provide a national collection of genetic resources to secure the biological diversity that underpins a sustainable U.S. agricultural economy.” Some of these collections are the target for high-density genotyping projects making them ideal populations for genome-wide association studies. However, the prohibitive cost of controlled field trials to measure novel phenotypes can limit their utility for genetics research. In this experiment, we used existing germplasm to find novel genotype-phenotype associations without the expensive overhead of independent field trials. Although this experiment is limited by the inability to grow plants in a common environment, the high heritability of ionomics traits (Baxter *et al.* 2014), as well as the stability of the ionome in stored tissue (Baxter *et al.* 2014), makes ionomic phenotyping an ideal test case for mining germplasm resources. To test the power of ionomics to find genetic factors underpinning elemental accumulation, we analyzed seeds from 1653 soybean [Glycine max (L.) Merr.] lines representing the diversity found in the USDA Soybean Germplasm Collection stored at Urbana, IL.

A core collection of 1685 accessions of the USDA Soybean Germplasm Collection represents a substantial amount of the genetic diversity in the entire collection. The core collection contains approximately 10% of the total number of introduced soybean accessions. The 1653 soybean lines used in this study comprised all of the 1685 accessions available when the research was started. For accessions in maturity groups 000 through VIII for which field evaluation data were available the core was selected using origin, qualitative and quantitative data. Accessions were divided in groups based on origin and then further subdivided based on maturity group, which classifies soybean accessions based on photoperiod and temperature response. A total of 81 strata were established. A multivariate proportional sampling strategy within each stratum was determined to be the optimal procedure for identifying a sample of accessions that best represents the diversity of the total collection. Field evaluation data were not available for accessions in maturity groups IX and X, but because these accessions are adapted to sub-tropical and tropical conditions and are likely to have unique genetic diversity, a sample of 10% of these accessions was added to the core collection based on multivariate analysis of the qualitative data. A full explanation of the development of the core collection can be found in Oliveira et al. (2010). The seeds available in the NPGS for this core collection come from grow-outs that span 12 years at three locations (Urbana, IL, Stoneville, MS, and Upala, Costa Rica) (Table 1). The selection of which lines to grow for line maintenances in a given year is independent of the strata used to select the core collection, making each growout year an independent experiment to look for loci controlling elemental accumulation. Additonally, analysis of the first two principal components from the SNP dataset shows no apparent bias between genetic architecture and growout (Supplemental Figure 1).

**Table 1.** Number of lines and markers in each GWAS dataset. There is no overlap between lines in the datasets. Markers are the number of segregating SNPs in each dataset, filtered for minor allele frequency > 0.05.

| Location | Growout Year | Lines | GWAS<br>Markers |
| --- | --- | --- | --- |
| Stoneville | 1999 | 104 | 33962 |
| Stoneville | 2004 | 121 | 34571 |
| Stoneville | 2006 | 59 | 35192 |
| Urbana | 2000 | 109 | 36432 |
| Urbana | 2001 | 69 | 36032 |
| Urbana | 2002 | 94 | 36151 |
| Urbana | 2003 | 147 | 35783 |
| Urbana | 2004 | 89 | 35490 |
| Urbana | 2005 | 87 | 35559 |
| Urbana | 2006 | 143 | 36065 |
| Urbana | 2007 | 98 | 36091 |
| Urbana | 2008 | 58 | 35432 |
| Urbana | 2009 | 102 | 36489 |
| Costa Rica | 9 years combined | 111 | 31479 |

**Figure 1.**
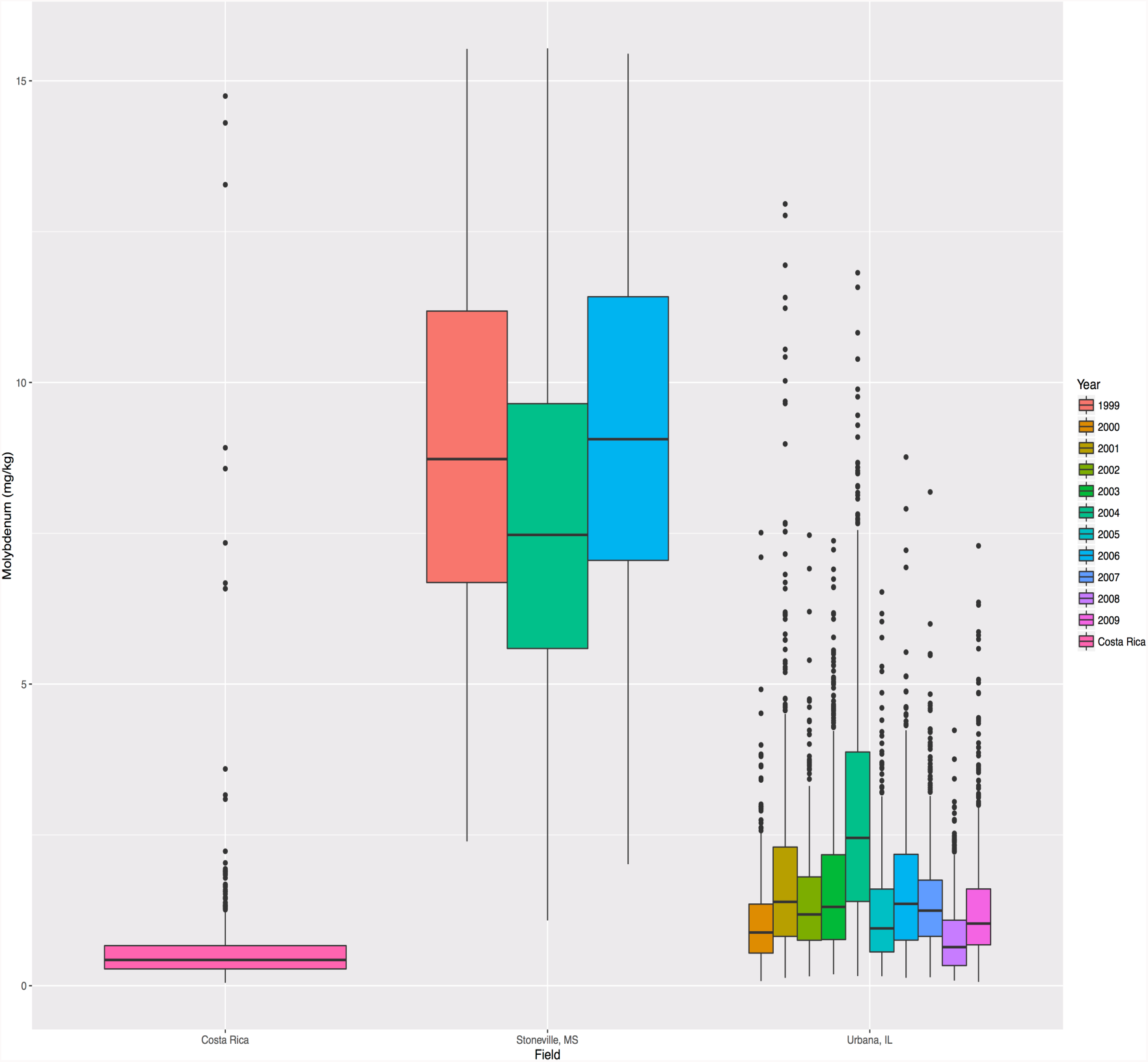
Molybdenum accumulation in single soybean seeds (mg/kg) across experimental grow-outs.

### Phenotypes

Using the elemental analysis pipeline described in Ziegler et al. (2013, see methods), we analyzed ~6 seeds from each line, measuring the levels of 20 elements in each seed (Supplemental Table 1). While 1653 lines were analyzed in total, 262 of these lines were from grow-outs containing fewer than 50 lines in the dataset. We excluded these lines from further analysis and all following analysis is based on the remaining 1391 lines (elemental profiles for excluded lines are included in the Supplemental Table 1). We performed an ANOVA significance test to assess whether there are significant environmental effects on the phenotypic data gathered from lines grown in separate locations and in separate years at the same location. Although a distinct set of lines were grown in each grow-out, lines were assigned to a grow-out without regard to population structure. As a result, we would expect, in the absence of environmental effects, phenotypic measurements to be similar. The ANOVA test indicates a significant location effect, and for Stoneville and Urbana, significant effects for growth year, for most elements measured (p<0.01 with Bonferroni correction, Table 2). This effect can also be seen in the phenotypic distribution (before transformation) for many of the traits (Figure 1 and Supplemental Figure 2). These results clearly demonstrate that most of the year growouts were unique environments, supporting their analysis as individual experiments. The lack of significant differences by year for many elements in Costa Rica (13 out of 21) may be indicative of a lack of statistical power due to the small number of lines grown per year. Because there were not enough lines in any one grow-out from Costa Rica for a GWAS analysis, the only way we were able to analyze the Costa Rica data was by combining data across all 10 years.

**Table 2.**
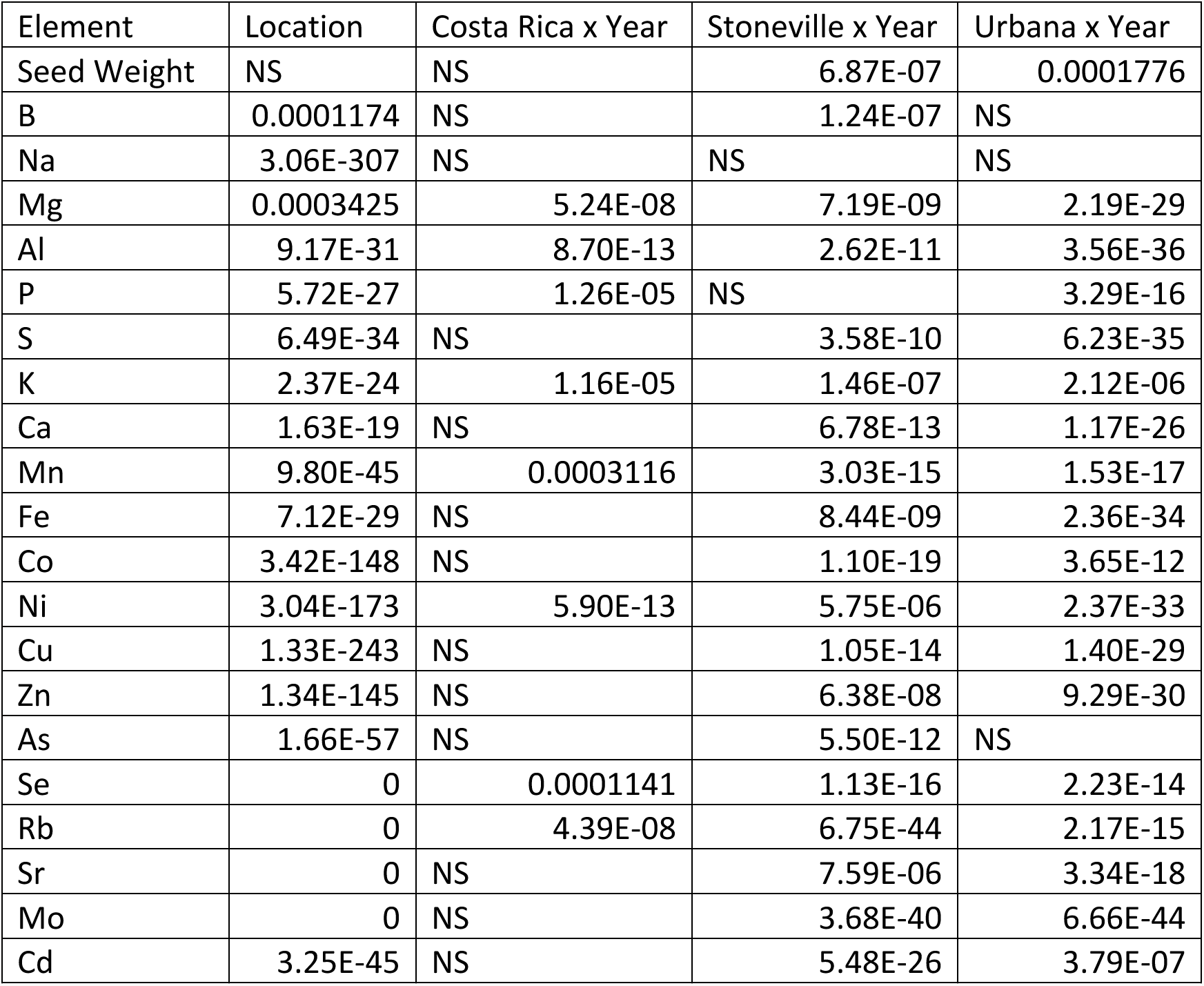
Analysis of grow out location and year effect on elemental accumulation. The *p*-value for each element from an ANOVA of a linear model with Location or Location x Year interaction. The significance cutoff was set at *p*< 0.01 with Bonferroni correction. NS=Not Significant

| Element | Location | Costa Rica x Year | Stoneville x Year | Urbana x Year |
| --- | --- | --- | --- | --- |
| Seed Weight | NS | NS | 6.87E-07 | 0.0001776 |
| B | 0.0001174 | NS | 1.24E-07 | NS |
| Na | 3.06E-307 | NS | NS | NS |
| Mg | 0.0003425 | 5.24E-08 | 7.19E-09 | 2.19E-29 |
| Al | 9.17E-31 | 8.70E-13 | 2.62E-11 | 3.56E-36 |
| P | 5.72E-27 | 1.26E-05 | NS | 3.29E-16 |
| S | 6.49E-34 | NS | 3.58E-10 | 6.23E-35 |
| K | 2.37E-24 | 1.16E-05 | 1.46E-07 | 2.12E-06 |
| Ca | 1.63E-19 | NS | 6.78E-13 | 1.17E-26 |
| Mn | 9.80E-45 | 0.0003116 | 3.03E-15 | 1.53E-17 |
| Fe | 7.12E-29 | NS | 8.44E-09 | 2.36E-34 |
| Co | 3.42E-148 | NS | 1.10E-19 | 3.65E-12 |
| Ni | 3.04E-173 | 5.90E-13 | 5.75E-06 | 2.37E-33 |
| Cu | 1.33E-243 | NS | 1.05E-14 | 1.40E-29 |
| Zn | 1.34E-145 | NS | 6.38E-08 | 9.29E-30 |
| As | 1.66E-57 | NS | 5.50E-12 | NS |
| Se | 0 | 0.0001141 | 1.13E-16 | 2.23E-14 |
| Rb | 0 | 4.39E-08 | 6.75E-44 | 2.17E-15 |
| Sr | 0 | NS | 7.59E-06 | 3.34E-18 |
| Mo | 0 | NS | 3.68E-40 | 6.66E-44 |
| Cd | 3.25E-45 | NS | 5.48E-26 | 3.79E-07 |

**Figure 2.**
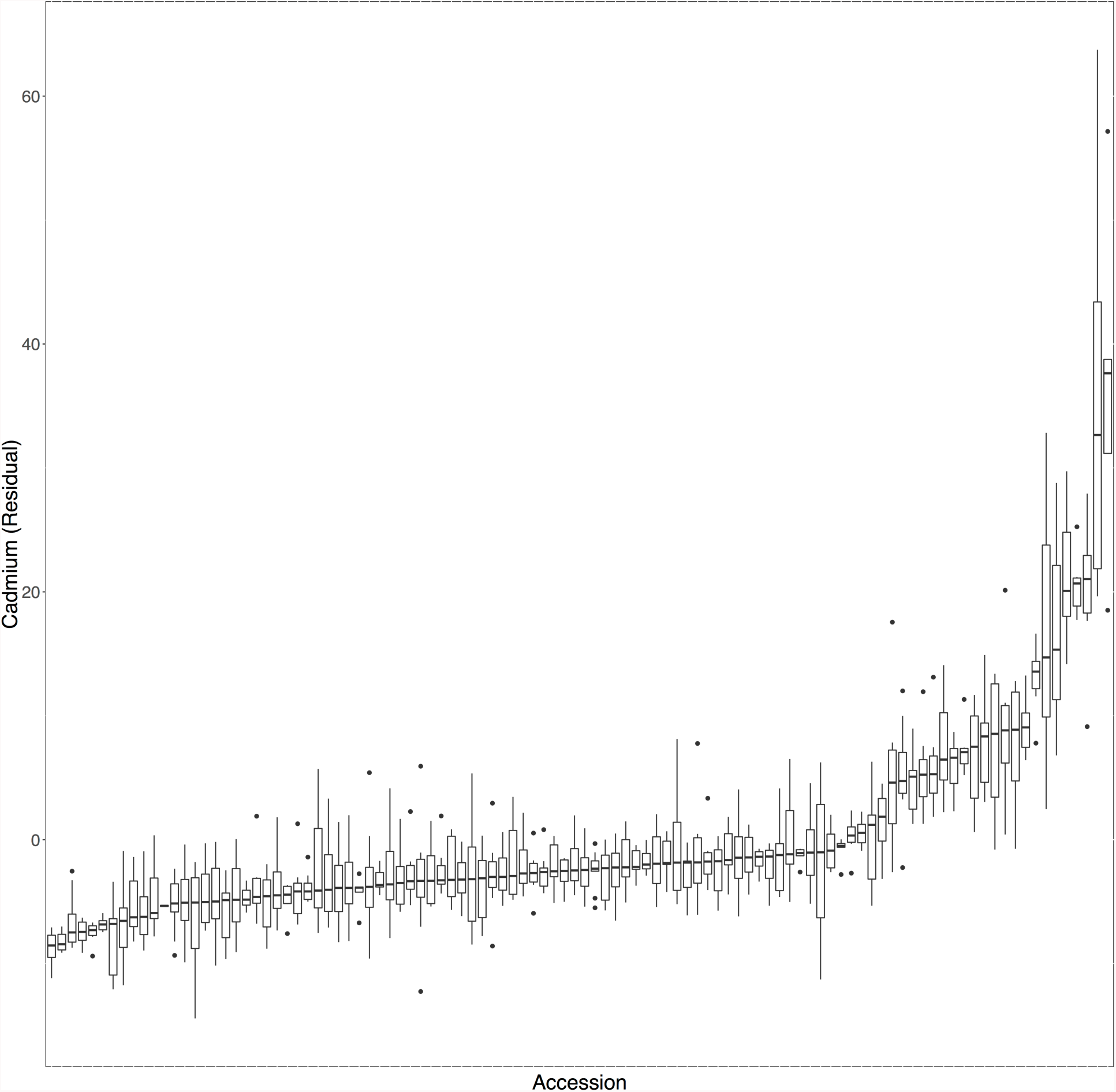
Distribution of Cadmium phenotype (linear model residuals, see Methods) in lines from a single growout: Stoneville, MS, 1999. Lines are ordered by median of between 2 and 8 seed replicates.

Comparison of elemental concentrations of replicate seeds from the same line in each grow-out does indicate the presence of a genotypic effect on elemental concentrations. Concentrations in seeds from the same line were usually more similar to each other than they were to the population as a whole (Figure 2 and Supplemental Figure 3).

The Box-Cox procedure (Box and Cox 1964) was used to estimate appropriate transformation functions for the phenotype data to meet the assumptions of GWAS for normally distributed dependent variables. The Box-Cox algorithm suggested that 138 of the 294 traits (14 environments × 21 phenotypes) needed no transformation and an additional 151 needed only minor transformations to control for the long-tail distributions often seen in concentration data (inverse, inverse square root, log, or square root) (Supplemental Table 2). Because most traits appear to only need minor transformations, for uniformity and ease of interpretation, all of the traits in which a transformation was recommended were transformed using a log transformation.

### Population Structure

The first two principal components obtained using the 36,340 polymorphic SNPs from the entire 1391 lines in the dataset explained 15% of the total SNP variance and the first 10 principal components explained 28% of the total variance. Variance explained by each PC drops rapidly after the first 10 PCs with 50% variance not reached until PC76. The first two principal components separate the population into groups roughly corresponding to each lines country of origin, with South Korean and Japanese accessions forming distinct clades while Chinese, Russian and other accessions form a much less cohesive block (Figure 3).

**Figure 3.**
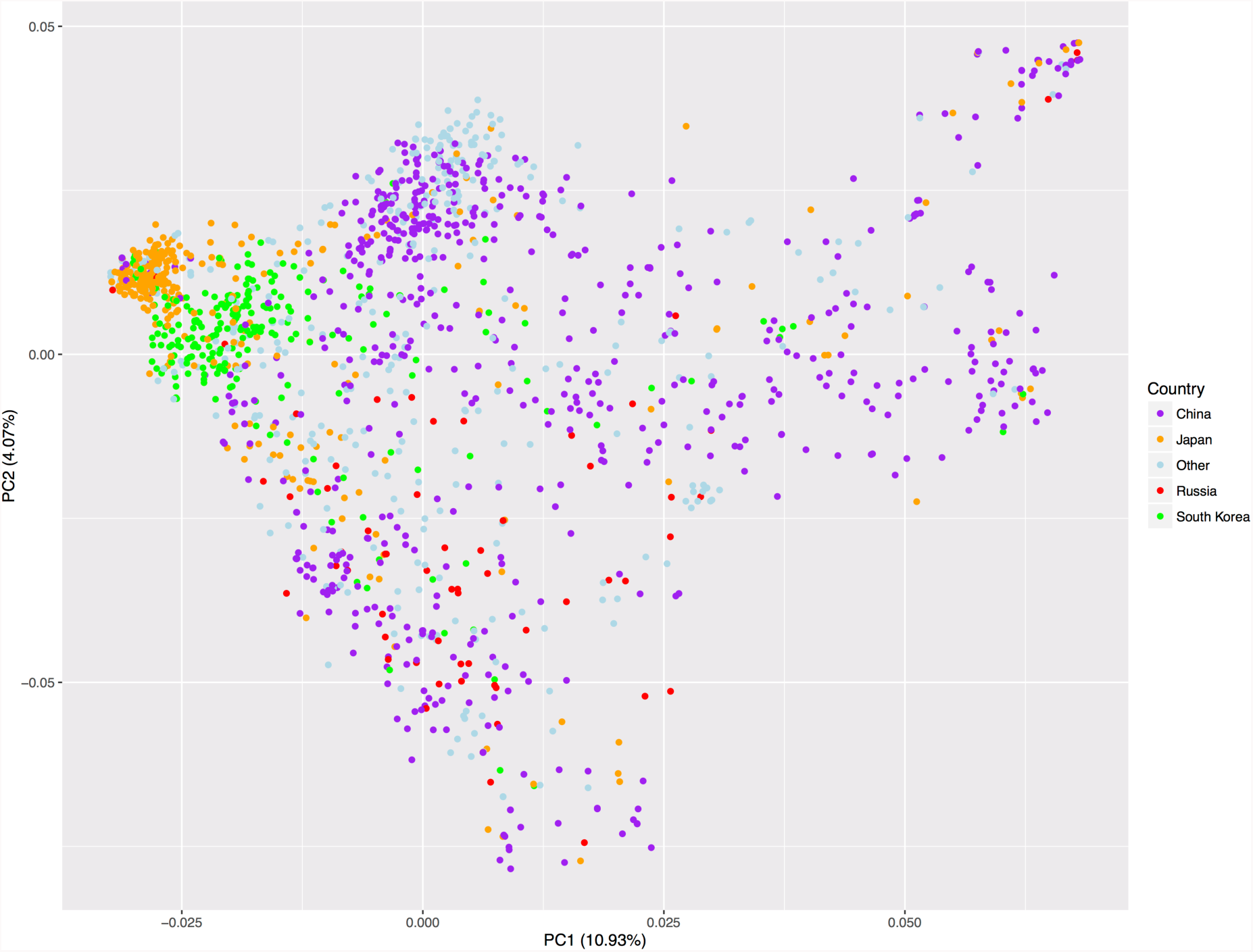
Principal component analysis of the genotypes of 1391 soybean lines. Colored by country of origin: China (532), Japan (267), South Korea (200), Russia (61), Other or unknown country of origin (331).

## MLMM GWAS

Using the SoySNP50k chip data (Song *et al.* 2013), we performed a GWAS study using a multi-locus mixed model (MLMM) to identify associated loci for each of 21 phenotypes (20 elements, seed weight) in 13 distinct grow-outs of diverse soybean lines and the Costa Rica dataset of grow-outs pooled across years (Table 1). The MLMM procedure starts with an EMMAX scan of all markers and then iteratively adds the markers with the highest association to the model and rescans. The MLMM procedure returns a list of cofactors that together describe the total estimated narrow-sense heritability of a given trait (which we will refer to as the all cofactor model). By definition, MLMM will create a model containing at least one cofactor for each trait. Of the models generated, 84 models met the stopping criteria after only one SNP was added to the model. The average model contained 11 SNPs, with no traits reaching the maximum 40 SNP model (e.g. not converging on a model describing all of the phenotypic variance). The largest model contained 29 SNPs, for iron in the 2009 Urbana grow-out. The 294 GWAS tests returned 1756 unique SNPs. While these most complex models likely contain factors that account for phenotypic variance merely by chance (e.g., false positives), many of these cofactors are likely real.

A simpler model, which includes only a subset of the total cofactors, can be selected using a model selection parameter (Segura *et al.* 2012). Segura et al. proposed two model selection criteria: the extended Bayesian information criterion (EBIC) and the multiple-Bonferroni criterion (mBonf) (Segura *et al.* 2012). Although both criteria produced generally similar results, we found the EBIC criteria to be less stringent than mBonf. Due to the relatively small sample size in many of our grow-outs, we have chosen the more inclusive EBIC criteria in an attempt to include more moderate effect loci in our model at the cost of a higher false positive rate. QQ-plots for both the null model, containing no cofactors, and the optimal EBIC model were generated to assess whether there were uncontrolled confounding effects in our model arising from cryptic relatedness and population structure. While there was some inflation of p-values in the null model, the MLMM procedure of iteratively including large-effect loci into the model successfully controls for this p-value inflation and the distribution of p-values in the EBIC models closely follows the expected null distribution except for the significantly associated loci (Figure 4 and Supplemental Figure 4).

**Figure 4.**
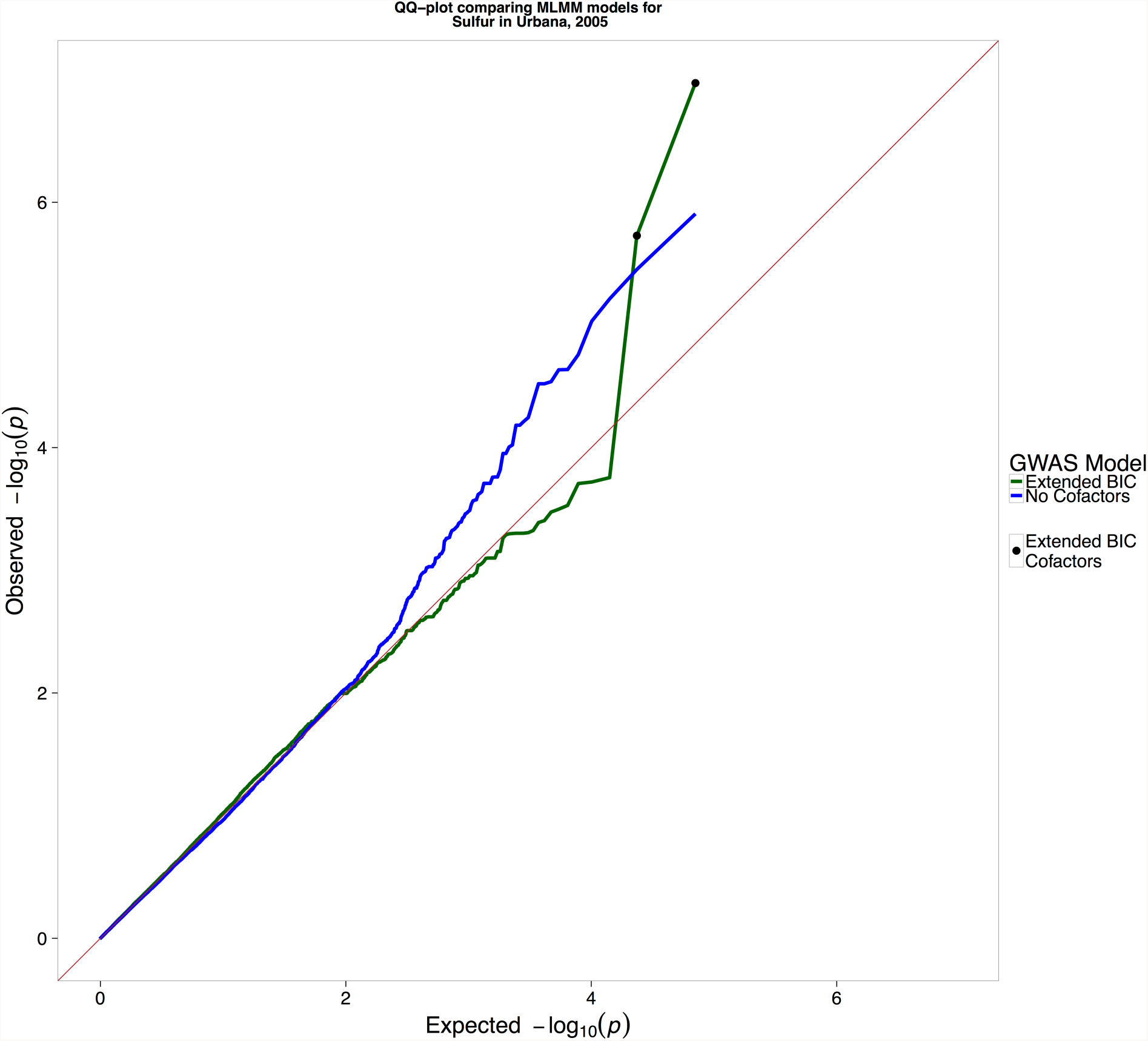
Quantile-quantile plot of the observed *p*-values against expected *p*-values from the GWAS analysis for sulfur accumulation. The MLMM algorithm includes cofactors that reduce inflation of *p*-values (green line). The model without cofactors indicates presence of *p*-value inflation (blue line). The expected distribution of *p*-values under the null hypothesis (red line).

The EBIC model selection method returned the MLMM model containing no cofactors for about half of the GWAS tests (164/294). The remaining 130 tests returned a total of 573 unique SNPs. When looking at the combined set of SNPs returned across all grow-outs, of the 21 phenotypes tested, at least one SNP was returned for each trait, with seed weight returning the most (96) and boron returning the least (6). Table 3 contains information about the number of cofactors returned in each model (EBIC and all) for each trait and Supplemental Table 3 contains the complete list of SNPs returned.

**Table 3.** Number of SNP cofactors returned by each GWAS experiment. Each cell contains the number of cofactors in the EBIC selected model and the all cofactor model, respectively. See methods for criteria for inclusion of a SNP in the EBIC or all cofactor model.

| Growout/<br>Element | Al | As | B | Ca | Cd | Co | Cu | Fe | K | Mg | Mn | Mo | Na | Ni | P | Rb | S | Seed<br>Weight | Se | Sr | Zn | Total |
| --- | --- | --- | --- | --- | --- | --- | --- | --- | --- | --- | --- | --- | --- | --- | --- | --- | --- | --- | --- | --- | --- | --- |
| 00U | 1/1 | 0/1 | 3/7 | 4/10 | 12/13 | 0/10 | 0/3 | 0/14 | 0/3 | 18/19 | 8/10 | 1/4 | 0/1 | 0/13 | 0/12 | 0/3 | 0/13 | 2/16 | 0/10 | 2/10 | 4/20 | 55/193 |
| 01U | 8/8 | 1/1 | 1/1 | 1/8 | 1/1 | 2/4 | 0/2 | 0/7 | 2/6 | 1/1 | 3/5 | 0/8 | 1/1 | 0/1 | 0/1 | 0/1 | 0/1 | 7/8 | 17/18 | 1/4 | 0/1 | 46/88 |
| 02U | 0/2 | 0/11 | 0/1 | 1/4 | 10/13 | 0/14 | 0/4 | 0/3 | 0/7 | 0/1 | 1/2 | 0/8 | 0/1 | 2/11 | 5/10 | 2/3 | 0/9 | 14/16 | 1/3 | 0/14 | 0/9 | 36/146 |
| 03U | 2/3 | 0/2 | 0/2 | 0/1 | 3/19 | 2/7 | 0/4 | 0/8 | 0/11 | 0/12 | 1/3 | 1/11 | 0/2 | 0/6 | 3/7 | 0/1 | 0/8 | 26/26 | 3/6 | 0/11 | 0/7 | 41/157 |
| 04S | 1/9 | 0/1 | 0/4 | 2/6 | 3/3 | 0/3 | 0/1 | 0/6 | 3/5 | 0/14 | 0/1 | 0/1 | 0/4 | 0/3 | 1/11 | 1/1 | 0/4 | 1/24 | 0/11 | 4/12 | 0/8 | 16/132 |
| 04U | 0/1 | 0/1 | 0/3 | 5/5 | 1/1 | 0/2 | 0/1 | 1/7 | 0/3 | 0/1 | 1/1 | 0/1 | 0/2 | 1/2 | 0/1 | 0/1 | 2/6 | 0/15 | 1/2 | 0/7 | 0/1 | 12/64 |
| 05U | 0/10 | 0/1 | 1/1 | 2/4 | 3/6 | 3/6 | 0/2 | 0/23 | 0/4 | 0/5 | 2/5 | 0/1 | 0/1 | 0/1 | 1/1 | 0/1 | 2/13 | 17/18 | 14/16 | 1/1 | 0/2 | 46/122 |
| 06S | 0/4 | 8/8 | 0/5 | 0/1 | 0/1 | 0/2 | 0/1 | 0/5 | 2/10 | 1/1 | 0/1 | 0/3 | 0/1 | 1/5 | 16/17 | 0/8 | 0/2 | 3/4 | 15/15 | 0/5 | 5/6 | 51/105 |
| 06U | 0/1 | 0/2 | 0/1 | 1/7 | 1/15 | 0/1 | 1/10 | 5/13 | 3/10 | 0/9 | 0/6 | 0/3 | 0/1 | 1/11 | 0/1 | 0/1 | 0/10 | 3/12 | 1/14 | 0/11 | 0/1 | 16/140 |
| 07U | 0/1 | 0/1 | 1/2 | 1/1 | 2/5 | 1/2 | 1/1 | 0/1 | 3/3 | 0/9 | 1/3 | 1/2 | 0/2 | 2/3 | 0/3 | 1/4 | 0/1 | 1/10 | 1/4 | 0/3 | 0/3 | 16/64 |
| 08U | 1/2 | 2/3 | 0/1 | 14/15 | 1/4 | 20/20 | 8/8 | 9/10 | 0/1 | 12/12 | 0/1 | 0/1 | 0/1 | 0/1 | 9/11 | 2/3 | 0/1 | 5/7 | 1/2 | 3/4 | 0/1 | 87/109 |
| 09U | 1/1 | 0/1 | 0/1 | 19/20 | 0/10 | 0/14 | 0/14 | 29/29 | 1/1 | 0/2 | 1/2 | 22/22 | 18/18 | 1/1 | 1/1 | 0/21 | 19/19 | 17/18 | 0/1 | 0/10 | 0/1 | 129/207 |
| 99S | 2/2 | 0/5 | 0/1 | 1/11 | 1/12 | 1/13 | 0/10 | 0/2 | 0/1 | 1/6 | 0/15 | 1/1 | 0/4 | 0/7 | 0/1 | 1/11 | 0/4 | 0/15 | 0/17 | 0/1 | 0/20 | 8/159 |
| CR | 0/11 | 0/1 | 0/3 | 0/8 | 4/7 | 0/11 | 0/1 | 2/3 | 7/8 | 3/11 | 1/7 | 7/9 | 0/3 | 0/4 | 0/9 | 0/8 | 0/9 | 0/12 | 0/1 | 2/13 | 0/12 | 26/151 |
| Total | 16/56 | 11/39 | 6/33 | 51/101 | 42/110 | 29/109 | 10/62 | 46/131 | 21/73 | 36/103 | 19/62 | 33/75 | 19/42 | 8/69 | 36/86 | 7/67 | 23/100 | 96/201 | 54/120 | 13/106 | 9/92 | 585/1837 |

Overall, despite a large number of tests for association (294), a relatively small number of SNPs were identified. Given the ability of the multi cofactor model to reduce the levels of spurious false positives, a large number of even the full model SNPS are likely to be real. However, given the large number of independent growouts and the partial independence of the elemental traits, we are able to apply more stringent criteria confidence in associations. Below, we list several sets of SNPs associated with elemental traits, ordered from ‘most confident’ to ‘lower confidence’. Since the likelihood of the same false associations being found more than once for the same trait in separate grow-outs with independent sets of lines is small, we looked for SNPs returned in multiple scans, which are likely to be real. Across these 130 experiments, 10 SNPs were returned more than once. Of these 10 SNPs, the exact same SNP was found for the same element in a different grow-out two times (ss715604985 and ss715605104, both for cadmium), different elements in the same grow-out once (ss715608340 for Ca and Sr), and different elements in different growouts 7 times (Table 4). The same element/multiple location and multiple element/same location SNPs constitute our highest confidence set for SNPs affecting the ionome, but likely greatly underestimate the useful information in the dataset.

**Table 4.** SNPs returned in the EBIC selected model in two or more grow-outs.

| Chromosome | Base Pair | Environment | Trait | logP | Model | Overlap Type |
| --- | --- | --- | --- | --- | --- | --- |
| 9 | 4612586 | 99S | Cd | 10.06 | EBIC | Same Element, Different Location |
| 9 | 4612586 | 04U | Cd | 5.39 | EBIC | Same Element, Different Location |
| 9 | 4991159 | 00U | Cd | 18.68 | EBIC | Same Element, Different Location |
| 9 | 4991159 | 02U | Cd | 18.95 | EBIC | Same Element, Different Location |
| 9 | 4991159 | 03U | Cd | 11.88 | EBIC | Same Element, Different Location |
| 9 | 4991159 | 06U | Cd | 6.77 | EBIC | Same Element, Different Location |
| 10 | 5863544 | 04S | Ca | 6.20 | EBIC | Different Element, Same Location |
| 10 | 5863544 | 04S | Sr | 7.68 | EBIC | Different Element, Same Location |
| 2 | 46468030 | 03U | Seed Weight | 11.73 | EBIC | Different Element, Different Location |
| 2 | 46468030 | 05U | Se | 29.18 | EBIC | Different Element, Different Location |
| 5 | 41315343 | 06S | Mg | 4.82 | EBIC | Different Element, Different Location |
| 5 | 41315343 | 09U | Mo | 4.58 | EBIC | Different Element, Different Location |
| 10 | 5179735 | 05U | S | 5.73 | EBIC | Different Element, Different Location |
| 10 | 5179735 | 06S | Ni | 7.36 | EBIC | Different Element, Different Location |
| 13 | 19554349 | 07U | Ni | 6.66 | EBIC | Different Element, Different Location |
| 13 | 19554349 | 09U | Ca | 18.06 | EBIC | Different Element, Different Location |
| 13 | 22047323 | 02U | Cd | 14.82 | EBIC | Different Element, Different Location |
| 13 | 22047323 | 06S | K | 5.59 | EBIC | Different Element, Different Location |
| 13 | 26504428 | 00U | Cd | 6.30 | EBIC | Different Element, Different Location |
| 13 | 26504428 | 03U | Seed Weight | 10.48 | EBIC | Different Element, Different Location |
| 19 | 84371 | 08U | Cu | 16.51 | EBIC | Different Element, Different Location |
| 19 | 84371 | 09U | Fe | 51.76 | EBIC | Different Element, Different Location |

Because each grow-out contains an independent set of lines, the set of SNPs tested differs between grow-outs depending upon the SNP minor allele frequency in each dataset. Additionally, common SNPs between growouts will still differ in allele frequency, which could result in neighboring SNPs, still in LD with the causal variant, being returned for different GWAS experiments. Therefore, looking for only exact overlaps between datasets may be overly restrictive. Soybean has been estimated to have a linkage disequilibrium (LD) decay distance of between 360Kbp in euchromatic regions and 9.6Mbp in heterochromatic regions (Hwang et al., 2014). To better search for overlaps between our datasets while also taking into account the large variability in LD range across the soybean genome, we grouped all of the SNPs returned across experiments by whether they are in LD with one another. Although many factors affect the ability to detect an association between a QTL and the actual causative loci, the minimal r^2^ for detection between the loci is generally estimated to be between 0.2 and 0.33 (Ardlie *et al.* 2002; Qanbari *et al.* 2010; Wallace *et al.* 2014) with a value of 0.2 previously being used to define LD range in the soybean genome (Hwang *et al.* 2014). Therefore, we defined an overlap between SNPs as whether a pair of SNPs has an r^2^ > 0.2. When this approach was applied to the all cofactors model, the same locus was returned for the same phenotype in different grow-outs 18 times, a different phenotype in the same grow-out 44 times and different phenotypes in different growouts 237 times (Supplemental Table 4). Often a SNP returned as significant in the EBIC model for one growout, will have a corresponding SNP in the all cofactor model of another growout, indicating that the signal is there in other populations, but at too weak a level to meet strict significance thresholds.

Another line of evidence that the SNPs identified are real is the co-location with candidate genes. Due to the large regions of linkage disequilibrium in the soybean genome, each of the 30,000 SNPs in our experiment is linked to dozens to hundreds of genes. Many plant processes, including root structure/function, water relations, and inter, intra and extra-cellular structures, can alter the elemental accumulation (Baxter *et al.* 2009; Tian *et al.* 2010; Chao *et al.* 2011, 2013; Barberon 2017). Each SNP is therefore likely to be associated with several plausible candidate genes. We looked under the SNPs of our overlap sets for strong candidates-those with orthologs associated directly with elemental phenotypes. Table 5 contains a list of SNPs found on or near candidate or already characterized genes. Many of the candidates are under SNPs associated with individual elements to which they or their orthologs were previously linked, or with chemically related elements (i.e Mn, Co, Cd with Fe or Se with S). The presence of these strong candidates under the detected SNPS supports the evidence from overlap that they are real associations.

**Table 5.** Returned SNPs overlapping candidate or already characterized genes. Bold font indicates lines returned in the more conservative EBIC model for at least one growout. SNP basepairs are mapped to soybean reference genome build Glyma1.1.

| 280<br>Chromosome | Base Pair (of most<br>significant SNP) | Environment(s) | Trait(s) | -logP (Of most<br>significant SNP) | Candidate Gene |
| --- | --- | --- | --- | --- | --- |
| **9** | **4991159** | 00U; 02U; 03U;<br>06U | Cd | 18.95 | HMA13; Glyma.09g055600 (Benitez et al., 2012); (Fang et al., 2016) |
| 2 | 43023030 | 99S; CR | Cd | 20.67 | Glyma.02g215700 is similar to At2-MMP which is induced during cadmium stress to leaves (Golldack et al., 2002) |
| **3** | **40883820** | 02U; 99S | Se | 21.15 | NRAMP metal transporter (Glyma.03g181400); Aluminum Sensitive 3 (ALS3; Glyma.03g175800) |
| 5 | 33737561 | CR; 09U | Ca | 36.24 | Multidrug resistance-associated protein 3 (MRP3, Glyma.05g145000); AtMRP5 implicated in Calcium homeostasis in Arabidopsis (Gaillard et al., 2008) |
| **14** | **47003645** | 06S; 03U | Co | 17.91 | ZIP metal ion transporter (Glyma.14g196200); Overlaps with a Zn and Rubidium (in all cofactor) |
| **15** | **410656** | 04S; 07U | Mn | 7.11 | CAX2 (Glyma.15g001600), implicated in Mn transport (Shigaki et al., 2002); NRAMP6 (Glyma.15g003500), Mn transport; MGT2 (Glyma.15g002700) and MGT4 (Glyma.15g005200), magnesium transport |
| **2** | **5555909** | 07U | Fe; Zn; P;<br>Cu | 6.91 | ATOX1 (Glyma.02g068700), Copper transport |
| **1** | **54551283** | 01U; CR; 00U;<br>04U | Al; Rb; Mo;<br>Co; K | 7.64 | ALMT (Glyma.01g223300), Aluminum activated malate transport, malate is a chelator for aluminum and critical in detoxification |
| 2 | 44460357 | 09U; 02U | Co; Ca | 10.96 | Heavy metal transport/detoxification (Glyma.02g222600, Glyma.02g222700); Potassium transporter 1 (Glyma.02g228500); Phosphate transporter 4;3 (Glyma.02g224200) |
| **3** | **5165511** | 09U; 06U | Fe; Mn | 36.05 | YSL6 (Glyma.03g040200); FPN1 ferroportin (Glyma.03g042500) |
| **7** | **5480577** | 06S; 06U | As; Ni | 22.46 | Heavy metal transport/detoxification (Glyma.07g065800); NRAMP2 (Glyma.07g058900) |
| **11** | **17367460** | 04U; 06U | Fe; Se | 21.13 | ABC Transporter (Glyma.11g194700, Glyma.11g196100) |
| **19** | **84371** | 08U; 09U | Cu; Fe | 51.76 | ATOX1 (Glyma.19g001000), Copper transport |
| **3** | **5455217** | 00U; 04U | Mg; Co | 7.45 | iron regulated 1 (Glyma.03g042500); iron regulated 2 (Glyma.03g042400); YSL6 (Glyma.03g040200) |
| **15** | **1222084** | 05U | Se | 29.64 | Sulphate Transporter (Glyma.15g014000) (El Kassis et al., 2007; Cabannes et al., 2011); Sulfite Transporter (Glyma.15g015600) |
| **9** | **4799335** | 06S | K | 4.31 | Potassium Transporter (Glyma.09g052700) |
| **7** | **5900018** | 06U | Fe | 5.07 | Overlap with IDC for FRO2 (Mamidi et al. 2014); Glyma.07g067700; Also Glyma.07g065800 a heavy metal detox |
| **9** | **4518093** | 09U | Mo | 17.96 | Molybdenum Cofactor sulfurase (Glyma.09g050100) |
| **9** | **3807440** | 09U | S | 31.98 | Glyma.09g045200 Heavy Metal Transport; Close to all cofactor selenium |
| 5 | 8074553 | 00U; 06S | Fe | 7.06 | Stabilizer of iron transporter (AGO10, PNH, ZLL; Glyma.05g011300), in IDC dataset (Mamidi et al. 2014) |
| 3 | 45338714 | 03U | Fe | 8.30 | NAS3; Glyma.03g231200; Overlaps IDC (Mamidi et al. 2014) |

### Verification of High and Low Sulfur and Phosphorus accumulating lines

To test whether the elemental accumulation of ionomic traits in the lines in our panel are intrinsic to the genetics of the lines or an artifact of the environmental and field conditions, we performed two experiments in which we selected the highest and lowest accumulating lines for sulfur and phosphorus and regrew the seeds in controlled field and greenhouse conditions. Eight lines, four with a high phosphorus phenotype and four with a low phosphorus phenotype were selected for regrowth in a field in Columbia, MO. Three of the four high phosphorus lines exhibited a high phosphorus phenotype in the regrow experiment, while the low phosphorus lines had phenotypes closer to the control line level (Figure 5 and Table 6). Broad-sense heritability for phosphorus between the GRIN growout concentrations and this experiment was 0.65 (Supplemental Table 5).

**Table 6.**
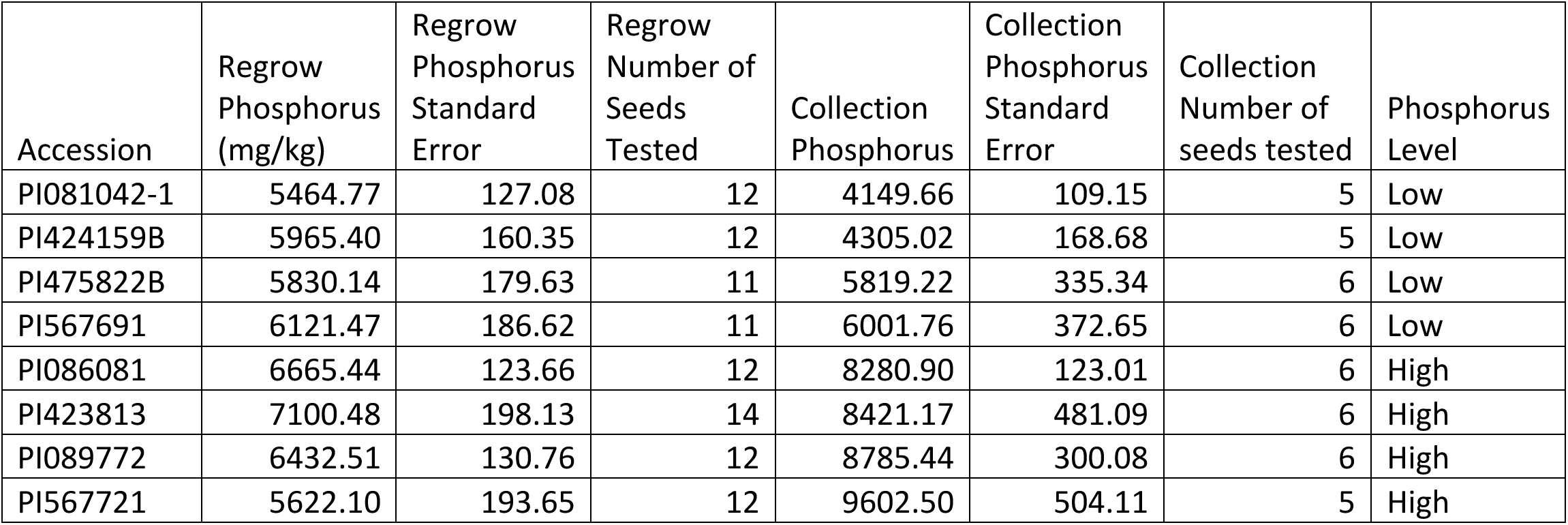
Accessions chosen for validation of phosphorus accumulation. High and low phosphorus accumulating lines were chosen to regrow to test the reproducibility of ionomic traits. Values listed in the table are mg Phosphorus/kg tissue.

In a separate experiment, 10 lines total, four low sulfur accumulating lines and six high sulfur accumulating lines were selected and regrown in both a field and greenhouse trial. In both the field and greenhouse experiment, all of the six high sulfur lines had a higher sulfur accumulation than the four low accumulating lines. Interestingly, the field grown varieties had a larger difference in sulfur accumulation between the high and low varieties (Figure 5 and Table 7). Although not selected for accumulation of other elements, there was also a correlation between measured values in the germplasm collection and the regrow set for many other elemental phenotypes tested (Supplemental Figures 5 and 6). Broad-sense heritability for sulfur between the GRIN growout concentrations, the greenhouse, and the field growouts was 0.64 (Supplemental Table 5).

**Table 7.** Accessions chosen for validation of sulfur accumulation. High and low sulfur accumulating lines were chosen to regrow to test the reproducibility of ionomic traits. Values listed in the table are mg sulfur/kg tissue.

| Accession | Regrow<br>Field<br>Sulfur<br>(mg/kg) | Regrow<br>Field<br>Standard<br>Error | Regrow<br>Field<br>Number of<br>Seeds<br>Tested | Regrow<br>Greenhouse<br>Sulfur<br>(mg/kg) | Regrow<br>Greenhouse<br>Standard<br>Error | Regrow<br>Greenhouse<br>Number of<br>Seeds<br>Tested | Collection<br>Sulfur<br>(mg/kg) | Collection<br>Sulfur<br>Standard<br>Error | Collection<br>Number of<br>seeds<br>tested | Sulfur<br>Level |
| --- | --- | --- | --- | --- | --- | --- | --- | --- | --- | --- |
| PI096322 | 3674.77 | 82.01 | 6 | 3303.99 | 86.76 | 6 | 2694.52 | 75.46 | 7 | Low |
| PI229327 | 3183.07 | 69.30 | 6 | NA | NA | NA | 2764.57 | 62.35 | 7 | Low |
| PI507411 | 3190.73 | 26.38 | 4 | 3126.35 | 84.73 | 6 | 2797.00 | 67.14 | 8 | Low |
| PI603599A | 3584.44 | 48.23 | 6 | 3075.94 | 114.71 | 8 | 2874.06 | 64.85 | 8 | Low |
| PI603162 | 4336.25 | 45.05 | 6 | 3703.22 | 70.82 | 6 | 3771.84 | 71.02 | 8 | High |
| PI339734 | 4856.20 | 158.22 | 6 | 4875.50 | 68.81 | 4 | 3774.48 | 21.99 | 2 | High |
| PI437377 | 4728.93 | 112.23 | 6 | 3413.30 | 82.30 | 6 | 3847.54 | 82.38 | 7 | High |
| PI603910B | 4301.96 | 64.81 | 5 | 4074.24 | 80.70 | 5 | 3925.33 | 71.42 | 8 | High |
| PI082278 | 4703.29 | 51.39 | 5 | 4265.62 | 99.98 | 6 | 3929.56 | 117.16 | 7 | High |
| PI424078 | NA | NA | NA | 4791.33 | 187.03 | 5 | 4245.06 | 78.57 | 5 | High |

**Figure 5.**
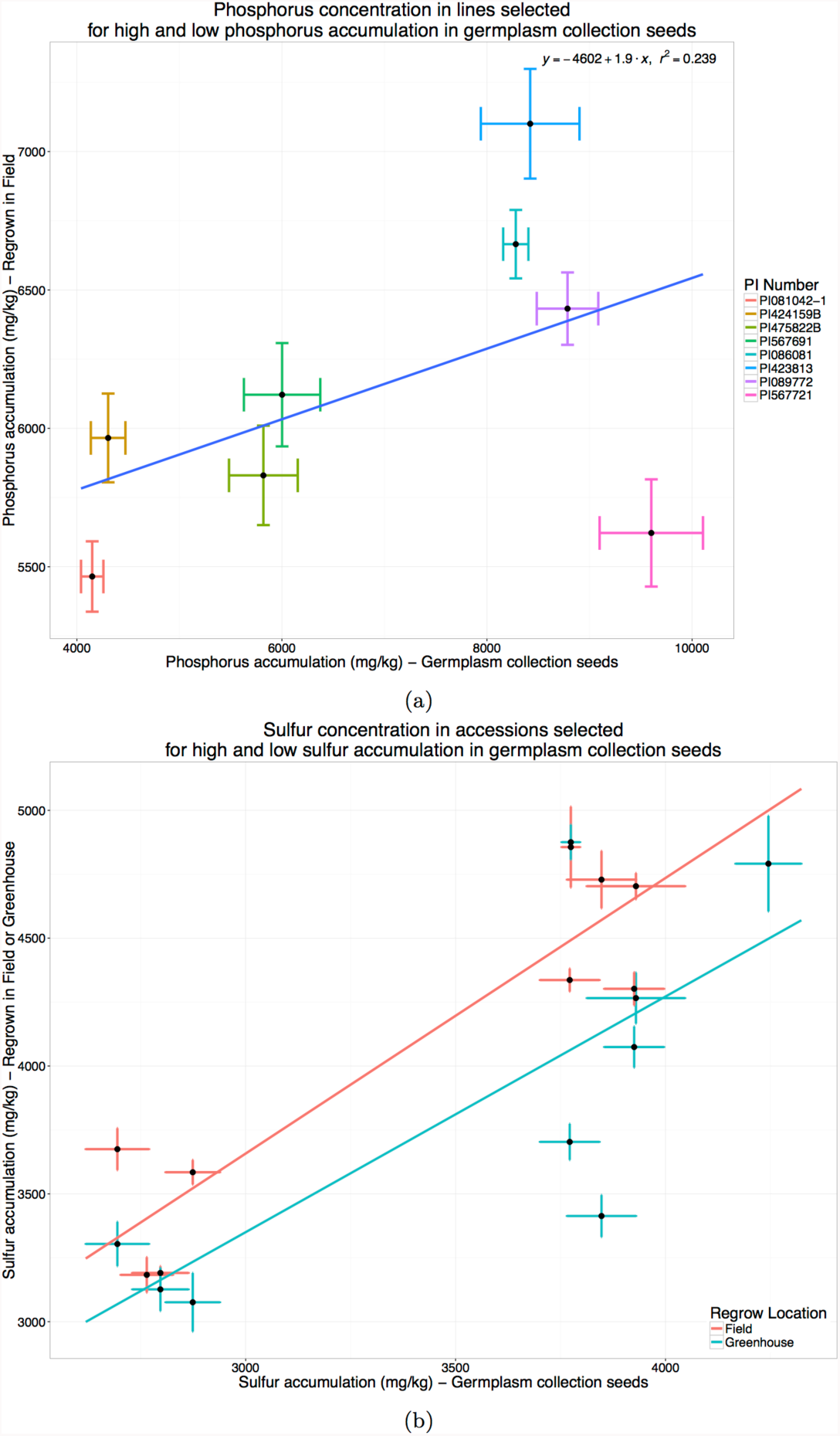
Confirmation grow out of high and low sulfur and phosphorus accumulating lines. A, Regrow versus original concentration of 8 lines selected for high and low phosphorus accumulation. Correlation between GRIN concentration and regrow was 0.24. B, Regrow versus original concentration of 10 lines selected for high and low sulfur accumulation, regrown in both greenhouse and field environments. Error bars indicate the standard error of the replicate seeds. Correlation (r^2^) between GRIN seed concentrations and the regrown high and low varieties grown in the greenhouse and in the fields were 0.61 and 0.84, respectively.

## Discussion

Analysis of ionomic traits has led to a deeper understanding of the complex regulatory system organisms use to maintain homeostasis of essential elements (Baxter *et al.* 2008; Baxter 2010; Atwell *et al.* 2010; Yu *et al.* 2012). To broaden our understanding of how genetic and environmental components affect the ionome, we have developed a high-throughput ionomic phenotyping system that can rapidly measure 20 ionomic traits and seed weight in agronomically important crops, such as soybean, maize, sorghum and cotton. To assess the utility of our phenotyping system for genome wide association studies in soybean, we measured the ionome of a diverse set of more than 1300 soybean lines, divided into 14 independent populations grown in three locations over the course of a decade. Coupled with a high-resolution genetic map (Song *et al.* 2013), we performed a genome wide association study using a multi-locus mixed model procedure (Segura *et al.* 2012). We were also able to show that lines selected from these experiments for extreme phenotypes of elemental accumulation were likely to display similar phenotypes in follow up experiments.

In spite of the limited number of lines in each grow-out, one of the strengths of this study is the number of distinct field replications. Although there was no overlap between lines for any of the 14 grow-outs, we found many genetic interactions that were robust across environments and genotypes. We report several different sets of SNPs corresponding to different levels of stringency in the individual experiments and the way we compared results between the experiments. These range from the 1756 SNPs from the full models, which likely contain several false positive associations, to the two SNPs that were returned in multiple experiments for the same element. Hundreds of SNPs in the total dataset are likely to be real due to their inclusion in a more conservative model or due to being found in several locations once LD is taken into account. Several of these mapped directly to what could be considered *a priori* candidate genes that have either already been characterized in soybean or are close orthologs of metal homeostasis proteins in *A. thaliana* and other species (Table 5). The discovery of orthologs of known *Arabidopsis* genes in soybean experiments highlights the value of studies in model organisms, where the genetics and growth habits are more amenable to large scale studies. Many more overlaps between different phenotypes found in different locations suggests genetic by environmental effect on which phenotype is affected by a causal locus. Many of the SNPs which overlap across environments are novel associations with no obvious gene candidates and are strong candidates for follow-up studies to determine their relationship to plant nutrient homeostasis.

The strongest element-loci association in our study was for the cadmium phenotype which is associated with a gene that codes for HMA13, a P_1B_-ATPase (HMA13; Glyma.09g055600) previously implicated in seed cadmium concentration in soybean (Benitez *et al.* 2012). A previous GWAS study on iron deficiency chlorosis found seven loci strongly associated with the disease phenotype (Mamidi *et al.* 2014). Our analysis returned 3 of the seven loci found in that study, all associated with seed Fe, including the two strongest associations from the IDC panel: a locus associated with nicotianamine synthase 3 (NAS3; Glyma.03g231200) and a locus associated with a stabilizer of iron transporter (AGO10; Glyma.05g011300). If gene discovery of small to medium effect loci is the goal of a study, using samples from germplasm banks may not be appropriate, but even with all the caveats about statistical power and gene by environment interactions, we found loci that had strong candidates for some elements. These results could be used to prioritize genes and lines for further characterization experiments.

## Conclusion

Using state-of-the-art association mapping techniques we were able to use the data we collected using our high-throughput ionomic phenotyping pipeline to identify both lines with extreme phenotypes and loci associated with elemental traits. Many of these associations were strong enough to occur across a diverse set of environmental conditions, while others were found in only one of the environments tested. While there are likely many more associations in our GWAS dataset that we haven't yet explored, this experiment serves as a proof of concept of using stored seed to perform GWAS on ionomic traits. While our efforts were focused on the identification of markers associated with elemental traits, the SNPs identified were associated with many *a priori* candidate genes. The use of seeds as the phenotyped tissue allows for the direct association of the consequences of allelic difference of SNPs and associated candidate genes with traits that affect the tissue with the most agronomic importance in soybeans. While planned experiments with more replication and higher numbers of lines will always have more power to identify genetic and environmental factors driving elemental accumulation in the seed, this study demonstrates the utility of leveraging available samples to screen germplasm.

## Materials and Methods

### Germplasm

A diverse panel of 1653 soybean accessions was selected from the core soybean collection of the USDA Soybean Germplasm Collection, as described in the results. Because the mission of NPGS is to maintain a viable collection of plant germplasm, the collections are periodically regrown to maintain viable seed. The size of the soybean germplasm collection necessitates that only a subset of the complete germplasm collection is grown-out each year. Furthermore, the diverse panel of accessions belongs to a variety of maturity groups and was grown-out in three separate locations: Stoneville, MS, Urbana, IL, and Upala, Costa Rica. The 1653 lines in the panel are, thus, broken into 13 distinct year and location sets, with no overlap of lines between years or locations (Table 1). The Costa Rica dataset had no individual years with enough lines (>50) to perform a successful association analysis. However, by creating three additional datasets by combining data from each location, regardless of year, we were able to analyze data from the Costa Rica grow-outs.

### Confirmation Growouts

Small plots of four low sulfur accumulating lines and six high sulfur accumulating lines were grown in Mexico silt loam soil at Bradford Research and Extension Center, Columbia, Missouri. Cultural practices were typical of those utilized for soybean production in the Midwest US. The same set of plants were also grown in environmentally controlled greenhouse in 6 liter pots containing PRO-MIX (Premier Horticulture, Quebec, Canada) medium amended with Osmocote Classic controlled release fertilizer (Scotts, OH). Greenhouse settings were 16 h day length with 30/18°C day/night temperatures.

Small plots of differential phosphorus lines were grown out in 2012 at South Farm Agricultural Research Center (Columbia, MO, Latitude 38.908189, Longitude −92.278693, Mexico silt loam soil) as single plots of 5 feet long with a 3 foot gap between rows and 30 inches between rows. Field conditions were typical of soybean production in the Midwest US, with NPK Fertilizer applied at rates appropriate according to soil analyses (10.6/50/75) and two pre-emergent herbicides were applied before planting: Authority First (Authority First Corp, Philadelphia, PA) applied at 6.45 oz/acre; and Stealth applied at 1 qt/ac (Loveland Products, Loveland, CO, USA). Post-emergent herbicides were also used: Ultra Blazer (UPI, King of Prussia, PA, USA) applied at 1.5pt/acre; Basagran (Arysta LifeScience North America, LLC, Cary, NC, USA) applied at 1.5pt/acre and Select Max (Valent Biosciences Corp., Libertyville, IL, USA) applied at 24 oz/acre. At maturity, plots were bulk harvested and threshed and a subsample was used for ICP-MS analysis.

### Ionomic Phenotyping by ICP-MS

Samples were phenotyped on two separate occasions for the elemental concentrations for B, Na, Mg, Al, P, S, K, Ca, Mn, Fe, Co, Ni, Cu, Zn, As, Se, Rb, Mo, and Cd following the analytical methods described in Ziegler et al. (2013). Seed weight is also recorded for each sample analyzed, so it was also included as a phenotype in our study.

A simple weight normalization procedure to correct measured sample concentrations for seed size was found to introduce artifacts, especially for elements whose concentration is at or near the method detection limit. This could either be due to a systematic over or under reporting of elemental concentrations by the ICP-MS procedure or a violation of the assumption that all elemental concentrations scale linearly with weight. We used an alternative method to normalize for seed weight following the method recently reported in Shakoor et al. (2016). A linear model was developed modeling unnormalized seed concentrations against seed weight and the analytical experiment the seed was run in. The residuals from this linear model were then extracted and used as the elemental phenotype. For each element, the phenotypic measurement was taken as the median of the elemental concentrations from the 2 or 8 seeds measured from each line (after outlier removal of measurements with a median absolute deviation of >10 where we had enough samples). To meet the normality assumptions required for GWAS, an analysis using the Box-Cox algorithm was used to determine an appropriate transformation for each trait (Box and Cox 1964). Since each grow-out has a distinct set of lines, which may result in different phenotypic distributions, transformations were performed separately for each element in each dataset listed in Table 1. Transformations were selected based upon the 95% confidence interval returned by the Box-Cox function implemented in the R package MASS (Box and Cox 1964; Venables *et al.* 2002).

### GWAS

All of the lines included in this analysis (and all of the annual accessions in the Soybean Germplasm Collection in 2010) have been genotyped using the SoySNP50K beadchip and are available at soybase.org (Song *et al.* 2013). Separate genotype files were generated for each grow-out that contain only the lines present in that grow-out. The genotype files were each filtered to remove SNPs with a minor allele frequency less than 0.05 and missing SNPs were imputed as the average allele for that SNP. The number of SNPs for each grow-out varied between 31,479 and 36,340. The final number of SNPs used for association mapping of each grow-out are listed in Table 1. SNPs were called using the Glyma1.1 reference genome. All SNP base pair locations reported are from a map to Glyma1.1.

Both kinship and structural components were included in the mixed model and were calculated using the filtered genotype matrix containing all 1391 lines found across all 13 grow-outs. The kinship matrix was calculated using the VanRaden method as implemented in GAPIT (VanRaden 2008; Lipka *et al.* 2012). To correct for population stratification a principal component analysis was performed. The first ten principal components were used as fixed effects in the mixed model.

Association mapping was performed using a multilocus mixed model (MLMM) approach that performs a stepwise mixed-model regression with forward inclusion and backward elimination of genotypic markers included as fixed effects (Segura *et al.* 2012). In this model forward steps are performed until the heritable variance estimate reaches 0 (indicating the current model includes covariates that explain all of the heritable phenotypic variance) or a maximum number of forward-inclusion steps have been performed, which we set at 40.

MLMM implements two model selection methods to determine the optimal mixed model from the set of step-wise models calculated: the extended Bayesian information criterion (EBIC, Chen and Chen 2008) and the multiple-Bonferroni criterion (mbonf, Segura *et al.* 2012). The EBIC model uses the Bayesian information criteria to select a model taking into account both number of SNPs in the analysis as well as number of cofactors in the model. In our analysis, the EBIC was usually less conservative (eg. selected larger models). A larger model likely increases the number of type 1 errors, but it is less likely to miss true associations. Because we are performing a further selection step comparing results across independent experiments, we used the EBIC models for further analysis. Additionally, we also analyzed the cofactors returned by the final forward inclusion model (maximum model), which includes either the maximum 40 cofactors or the total number of cofactors needed to explain the estimated heritability.

SNPs included as cofactors in either the EBIC model or the maximum model were compared across GWAS experiments. SNPs were determined to overlap with a neighboring SNP if it had an r^2^ LD of >0.2.

### Calculation of Linkage Disequilibrium

Linkage disequilibrium, expressed as a correlation coefficient between markers (r^2^), was calculated using the filtered SNP data set containing all 1391 lines from the experiment and the LD function of the ‘genetics’ R package (Warnes *et al.* 2013).

### Germplasm and Data Availability

Lines used can be found at the USDA Soybean Germplasm Center. All scripts and data used can be found at www.ionomicshub.org and https://github.com/baxterlab/SoyIonomicsGWAS.

## Figure/Table Legends

Supplemental Figure 1. Principal component analysis of the genotypes of 1391 soybean lines. Colored by GRIN growout.

Supplemental Figure 2. Elemental accumulation in soybean seeds across experimental grow-outs.

Supplemental Figure 3. Distribution of all elemental phenotypes in all grow-outs. Lines are ordered by the median of between 2 and 8 seed replicates.

Supplemental Figure 4. QQ-plots for all GWAS experiments performed.

Supplemental Figure 5. Regrow versus original concentration for all phenotypes in the phosphorus selection experiment.

Supplemental Figure 6. Regrow versus original concentration for all phenotypes in the sulfur selection experiment.

Supplemental Table 1. Raw ionomics data and phenotypes after transformation for GWAS for all lines in the experiment.

Supplemental Table 2. Box-Cox suggested transformations for ionomics phenotypes.

Supplemental Table 3. All SNPs returned in either ‘All Cofactor’, ‘EBIC’, or ‘Multiple Bonferroni’ models for all GWAS experiments.

Supplemental Table 4. SNPs returned in two or more grow-outs based on Linkage Disequilibrium calculation.

Supplemental Table 5. Broad-sense heritabilities calculated for ionomic traits in the sulfur and phosphorus confirmation experiments.

